# Possible Synergistic Modulation of Hormonal Balance and Ovarian Structure by Clomiphene–Letrozole Co-Administration in a Rodent Model of Hyperandrogenism

**DOI:** 10.1101/2025.11.02.686116

**Authors:** Olakunle J. Onaolapo, Olufemi O. Aworinde, Kehinde J. Olufemi-Aworinde, Adejoke Y. Onaolapo

## Abstract

Infertility, defined as the inability to achieve pregnancy after 12 months of unprotected intercourse, is a major reproductive health concern. In females, hyperandrogenism often contributes to polycystic ovarian syndrome (PCOS), a leading cause of infertility. Although clomiphene and letrozole are widely used as ovulation inducers, their individual efficacy is limited, and the potential benefit of combined therapy remains unclear. This study investigated the effects of clomiphene–letrozole co-administration on gonadotrophic hormones, inflammatory cytokines, antioxidant status, and ovarian histomorphology in a rat model of hyperandrogenism. Thirty female Wistar rats were randomised into five groups (n=6). Group A received saline, while groups B–E were administered testosterone enanthate (10 mg/kg, subcutaneous injection) for 35 days to induce PCOS. Group B served as PCOS control, while groups C, D, and E were additionally treated with clomiphene (100 µg/kg), letrozole (5 mg/kg), or their combination, respectively, for 10 days from day 36. Hormonal assays, cytokine profiling, antioxidant measurements, and ovarian histology were performed. Results showed that the co-administration significantly reduced body and ovary weights, lowered glucose levels, and improved oestradiol and follicle-stimulating hormone profiles compared with PCOS controls. Combination therapy also enhanced antioxidant capacity, reduced lipid peroxidation, and modulated inflammatory cytokines by lowering IL-1β and TNF-α while elevating IL-10. Histological evaluation revealed cystic follicles with basement membrane thickening in the PCOS control, consistent with ovarian hyperstimulation; and a reversal with clomiphene and or letrozole treatment. In conclusion, clomiphene–letrozole co-administration demonstrated superior benefits over monotherapy in modulating endocrine, oxidative, and inflammatory parameters, suggesting a potential therapeutic advantage in ovulation induction for PCOS-related infertility.

## 1.0 Introduction

Polycystic ovary syndrome (PCOS) is one of the most common causes of female infertility, accounting for approximately 55%–70% of cases, b; due to chronic anovulation [1–3]. Globally, its prevalence ranges between 5% and 20% [1–3]. Over the past few decades, advances in the understanding of PCOS pathophysiology have redefined it as both an endocrine and metabolic disorder affecting women of reproductive age [4,5]. Although its precise aetiology remains unclear, environmental factors such as nutrition and obesity are strongly implicated [3]. Clinically, PCOS is characterised by anovulation or oligo-ovulation, alongside hypothalamic– pituitary–ovarian axis dysfunction. Unlike other ovarian causes of infertility, the ovaries in PCOS were believed not to be structurally-defective but functionally dysregulated.

Current concepts suggest that PCOS evolves through a vicious cycle of hyperandrogenism, central obesity, insulin resistance, and hypothalamic–pituitary dysfunction, culminating in anovulation and infertility [6]. Therapeutic approaches are thus directed at alleviating key symptoms, including hyperandrogenism, menstrual irregularity, and infertility [5]. Interventions include lifestyle modification (diet, exercise, behavioural changes), surgical options (laparoscopic ovarian drilling, wedge resection), and pharmacotherapy. Pharmacological treatments encompass insulin sensitisers such as metformin for metabolic complications, and ovulation-inducing agents such as clomiphene and letrozole [3, 7, 8]. Clomiphene citrate, a selective oestrogen receptor modulator, exerts anti-oestrogenic effects at the hypothalamic level, whereas letrozole, a third-generation aromatase inhibitor, prevents androgen-to-estrogen conversion in peripheral, ovarian, and central tissues [3, 7, 8]. Since its introduction two decades ago, letrozole has been shown in multiple studies to achieve superior live birth rates compared with clomiphene in women with PCOS [3, 8, 9–11].

More recently, the potential benefits of clomiphene–letrozole co-administration have been explored, particularly in women resistant to either drug [10, 12, 13]. While some studies support the combination as an effective first-line or rescue therapy in severe PCOS, others report only marginal improvements in pregnancy rates, without statistically-significant advantages over letrozole monotherapy [14]. While PCOS is a leading cause of anovulatory infertility, affecting 5–20% of women worldwide and accounting for over half of infertility cases and being associated with a high burden; current treatment options remain suboptimal. Although clomiphene citrate and letrozole are widely used first-line ovulation inducers; their individual efficacy is limited, with variable live birth rates and cases of drug resistance. Emerging studies have suggested that combining clomiphene and letrozole may enhance ovulation and pregnancy outcomes, particularly in resistant cases, but findings remain inconsistent and inconclusive. Furthermore, the mechanistic effects of this combination on endocrine regulation, oxidative balance, inflammatory cytokines, and ovarian morphology are poorly-understood. These knowledge gaps necessitate experimental co-administration in PCOS. This study examined the effects of clomiphene, letrozole, and their co-administration on gonadotrophic hormone profiles in a rodent model of hyperandrogenism. It would also determine the impact of these treatments on ovarian oxidative stress markers, total antioxidant capacity, pro-inflammatory (IL-1β, TNF-α) and anti-inflammatory (IL-10) cytokine levels; while examining the histomorphological changes in the ovaries following treatment with clomiphene, letrozole, and their co-administration. Finally, it compared the overall therapeutic benefits of clomiphene–letrozole co-administration relative to monotherapy, in reversing PCOS-associated alterations.

### 2.0 Materials and Methods

### 2.1 Chemicals and drugs

Letrozole 2.5 mg tablets (Novartis Pharma), Clomiphene citrate (Clomid^®^) 50 mg tablets Testosterone enanthate, Corn oil.

### 2.2 Animals

Healthy prepubertal female Wistar rats used in this study were obtained from the animal house of the Ladoke Akintola University of Technology Ogbomoso, Oyo State, Nigeria. Rats were housed in wooden cages measuring 20 x 10 x 12 inches in temperature-controlled (22.5°C ±2.5°C) quarters with lights on at 7.00 am. Rats were allowed free access to food and water. All procedures were conducted in accordance with the approved protocols of the Ladoke Akintola University of Technology and within the provisions for animal care and use prescribed in the scientific procedures on living animals, European Council Directive (EU2010/63).

### 2.3 Diet

All animals were fed commercially available standard rodent chow (29% protein, 11% fat, 58% carbohydrate) throughout the study.

### 2.4 Experimental methodology

Thirty weaned (21 day old) female rats weighing 60—70 g each were randomly assigned into five groups of six (n=6) animals each. Group A served as normal control and were administered vehicle (normal saline) orally at 10 ml/kg. Group B served as polycystic ovarian syndrome (PCOS) control and were administered testosterone enanthate subcutaneously at 1mg/100 g body weight [15, 16]. Rats in groups C and D were administered clomiphene (at 100 microgram/kg body weight) or letrozole at 5mg/g body weight respectively. Animals in group E were co-administered clomiphene and letrozole. Testosterone was administered daily for 35 days, while beginning on day 36, clomiphene, letrozole or distilled water was administered daily for 10 days [16]. Feed intake and body weight were measured using a weighing scale. At the end of the experimental period, animals were euthanised by cervical dislocation and blood was taken via an intracardiac puncture for the estimation of hormonal levels {oestrogen, progesterone, luteinizing hormone (LH), follicle stimulating hormone (FSH)}, inflammatory cytokines (interleukin 10, interleukin 1β and TNF-α), lipid peroxidation (measured as malondialdehyde concentration) and antioxidant status (Total antioxidant capacity). The ovaries were removed, observed grossly, weighed and fixed in 10 % neutral buffered formalin. Sections of the ovaries were then processed for paraffin-embedding, cut at 5 µm and processed for general histological study.

### 2.5 Assessment of body weight and food intake

Daily estimation of feed Intake and weekly measurement of body weight of animals in all groups was done using an electronic weighing scale (Mettler Toledo Type BD6000, Switzerland) as previously described [17, 18]. Percentage change in body weight or feed intake was calculated for each animal using the equation below following which results for all animals were computed to find the statistical mean

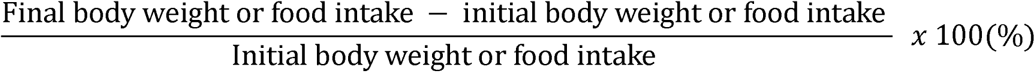

### 2.6 Assessment of relative weight of the ovaries

The ovaries were carefully dissected, weighed and washed in cold PBS following which it was blotted dry on a filter paper as previously described [19]. The weight of the ovary and body weight at sacrifice were taken and relative ovarian weight calculated using the formula below

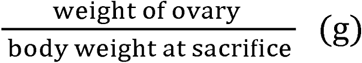

### 2.7 Histopathological study

Fixed samples of the ovary were embedded in paraffin and sectioned (at 5μm thickness). Sections of the ovarian tissue were then dewaxed and rehydrated following which they were mounted on slides and stained with hematoxylin–eosin (H&E) for general histological study while histopathological assessment of the connective tissue of the ovaries was done using the Verhoeff’s Van Gieson stain [20].

### 2.8 Biochemical assays

#### 2.8.1 Estimation of MDA content (Lipid peroxidation)

Lipid peroxidation level was measured as malondialdehyde content as described previously [21, 22]. Change in colour was measured using a spectrophotometer at 532 nm.

#### 2.7.1 Antioxidant activity

Superoxide dismutase activity was determined using commercially available assay kit. Colour changes were measured at an absorbance of 560 nm as described previously described [23,24]. The activity of SOD was expressed as units/ml.

Total antioxidant capacity was based on the principle of the Trolox equivalent antioxidant capacity and assayed following previously described protocols [25, 26].

#### 2.7.2 Glucose estimation

Glucose was measured using the glucose oxidase method as previously-described [27, 28].

#### 2.7.3 Lipid profile

Total cholesterol, triglycerides, HDL-C, and LDL-C in serum were analysed using commercially-available kits (RANDOX Laboratories, Crumlin, Co. Antrim, UK) as previously described [29, 30].

#### 2.7.4 Tumour necrosis factor-α, Interleukin (IL) −10, and interleukin 1β

Tumour necrosis factor-α and interleukin (IL)-10 were measured using enzyme-linked immunosorbent assay (ELISA) techniques with commercially-available kits (Enzo Life Sciences Inc. NY, USA) designed to measure the ‘total’ (bound and unbound) amount of the respective cytokines as previously described ([25, 26]. Interleukin-1 β level were assayed using enzyme-linked immunosorbent assay (ELISA) techniques with commercially-available kits (Enzo Life Sciences Inc. NY, USA) following the instructions of the manufacturers.

#### 2.7.5 Oestradiol, progesterone, Luteinizing and Follicle stimulating hormone assay

Oestradiol, progesterone, follicle stimulating hormone (FSH), and luteinizing hormone (LH) were measured using commercially-available assay kit (ELISA) kit (ABNOVA Taipei, 11493 Taiwan).

### 2.8 Photomicrography

Sections of the ovaries were examined microscopically using a Sellon-Olympus trinocular microscope (XSZ-107E, China) with a digital camera (Canon Powershot 2500), and photomicrographs taken. Histopathological changes were assessed by a technician that was blinded to the groupings. The thickness of the epithelial layer of the ovaries was also measured.

### 2.9 Statistical analysis

Data were analysed with Chris Rorden’s ANOVA for windows (version 0.98). Data analysis was by One-way analysis of variance (ANOVA) and post-hoc test (Tukey HSD) was used for within and between-group comparisons. Results were expressed as mean ± S.E.M. and p < 0.05 was taken as the accepted level of significant difference from control.

### 4.0 Result

### 4.1 Effect of clomiphene and letrozole on body weight

Figure 1 shows the effect of clomiphene and letrozole on weekly body weight (upper panel) and relative change in body weight (lower panel) in testosterone-treated female rats. There was a significant (p<0.001] increase in weekly body weight in the group administered testosterone alone (PCOS control), clomiphene (CLOM), Letrozole (LETR) and CLOM/LETR compared to control. Animals in the groups administered clomiphene and letrozole had the slowest trend in weekly body weight. Relative change in weight (percentage weight gain increased significantly (p<0.001) with PCOS and LETR, and decreased with CLOM and CLOM/LETR compared to control. Compared to PCOS, percentage weight gain was significantly decreased in the groups administered CLOM and CLOM/LETR.

**Figure 1:**
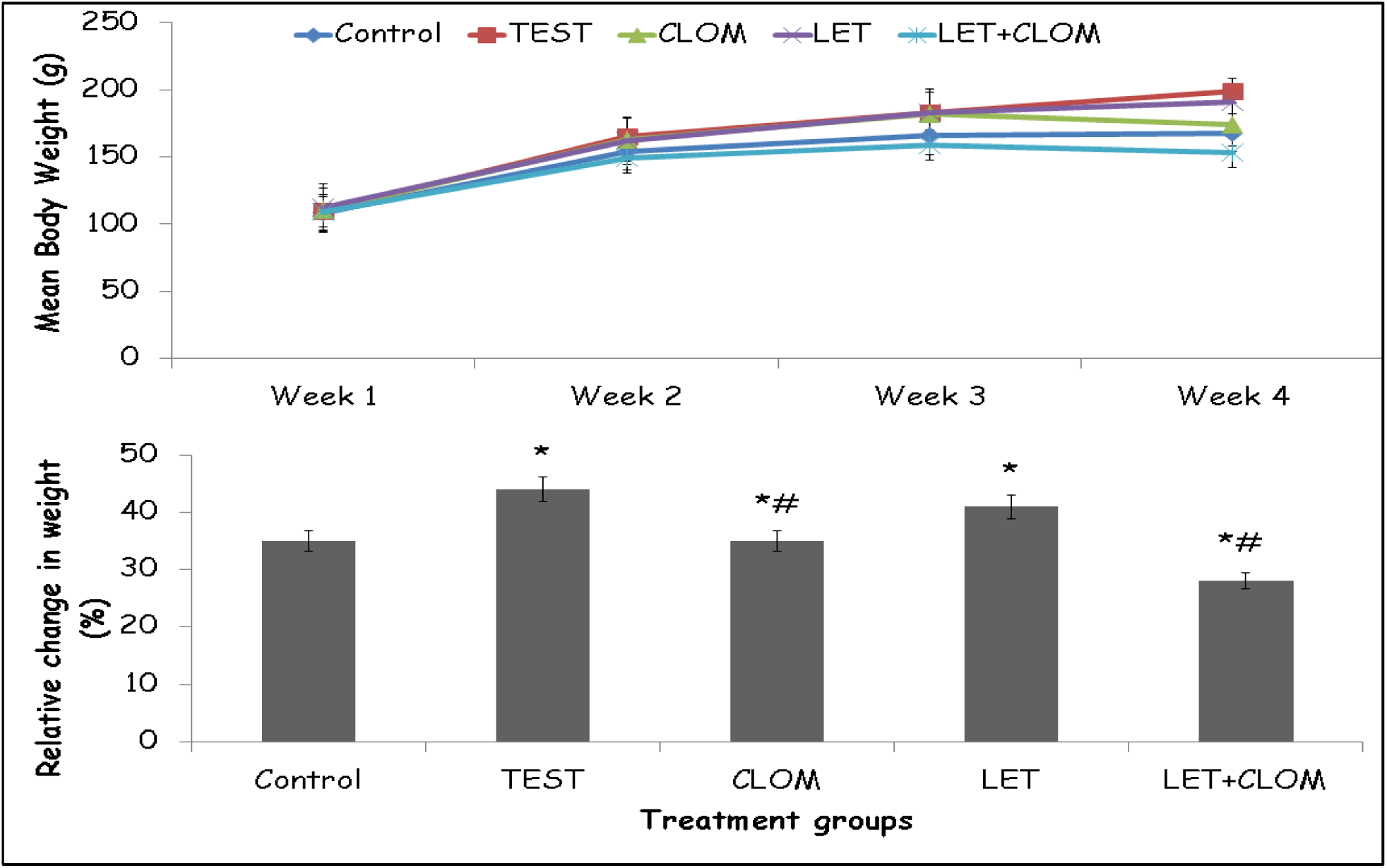
Effect of clomiphene and letrozole on weekly body weight (upper panel) and percentage change in weight (lower panel) in testosterone treated female rats. Each bar represents Mean ± S.E.M, number of mice per treatment group =6. PCOS: Polycystic ovarian syndrome, CLOM: Clomiphene, LETR: Letrozole.

### 4.2 Effect of clomiphene and letrozole on feed intake

Figure 2 shows the effect of clomiphene and letrozole on weekly feed intake (upper panel) and relative change in food intake (lower panel) in testosterone treated female rats. There was a significant (p<0.001] increase in weekly food intake with CLOM, LETR; and a decrease with CLOM/LETR compared to control and PCOS control respectively. Animals in the groups administered clomiphene and letrozole consumed the least food weekly throughout the experimental period. Relative change in food intake (Percentage change in food intake) increased significantly (p<0.001) with CLOM, LETR and decreased with CLOM/LETR compared to control. Compared to PCOS, percentage change in food intake was significantly increased with CLOM and LETR, and decreased with CLOM/LETR.

**Figure 2:**
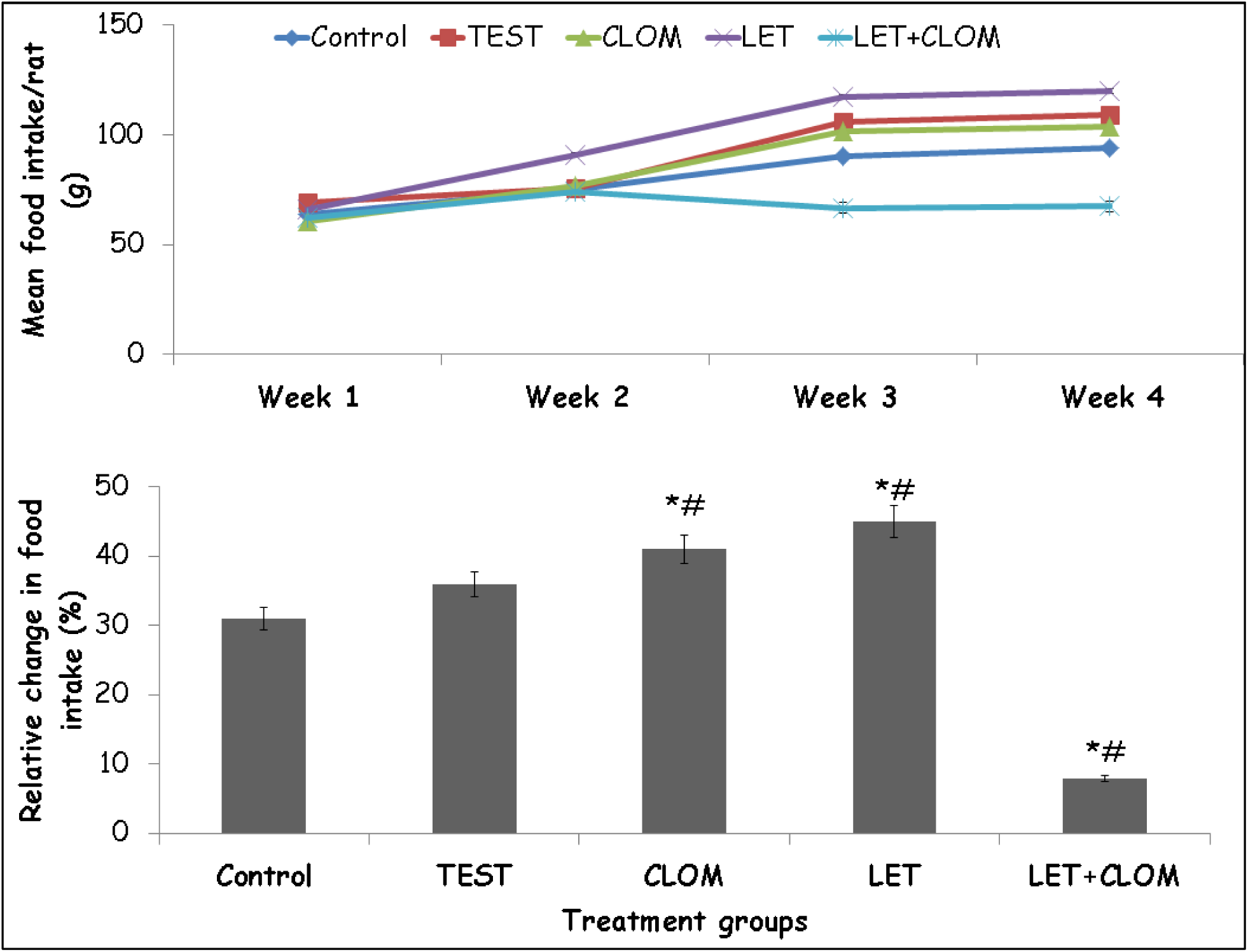
Effect of clomiphene and letrozole on weekly food intake (upper panel) and percentage change in food intake (lower panel) in testosterone treated female rats. Each bar represents Mean ± S.E.M, number of mice per treatment group =6. PCOS: Polycystic ovarian syndrome, CLOM: Clomiphene, LETR: Letrozole.

### 4.3 Effect of clomiphene and letrozole on ovary weight

Figure 3 shows the effect of clomiphene and letrozole on ovary weight (upper panel) and relative weight of the ovary (lower panel). Ovary weight increased significantly with PCOS and CLOM compared to control. Compared to PCOS control, ovary weight decreased significantly with CLOM, LET and CLOM/LET. Relative ovary weight increased significantly with PCOS and CLOM compared with control. Compared to PCOS control, relative weight of the ovary decreased with CLOM, LET and CLOM/LET.

**Figure 3:**
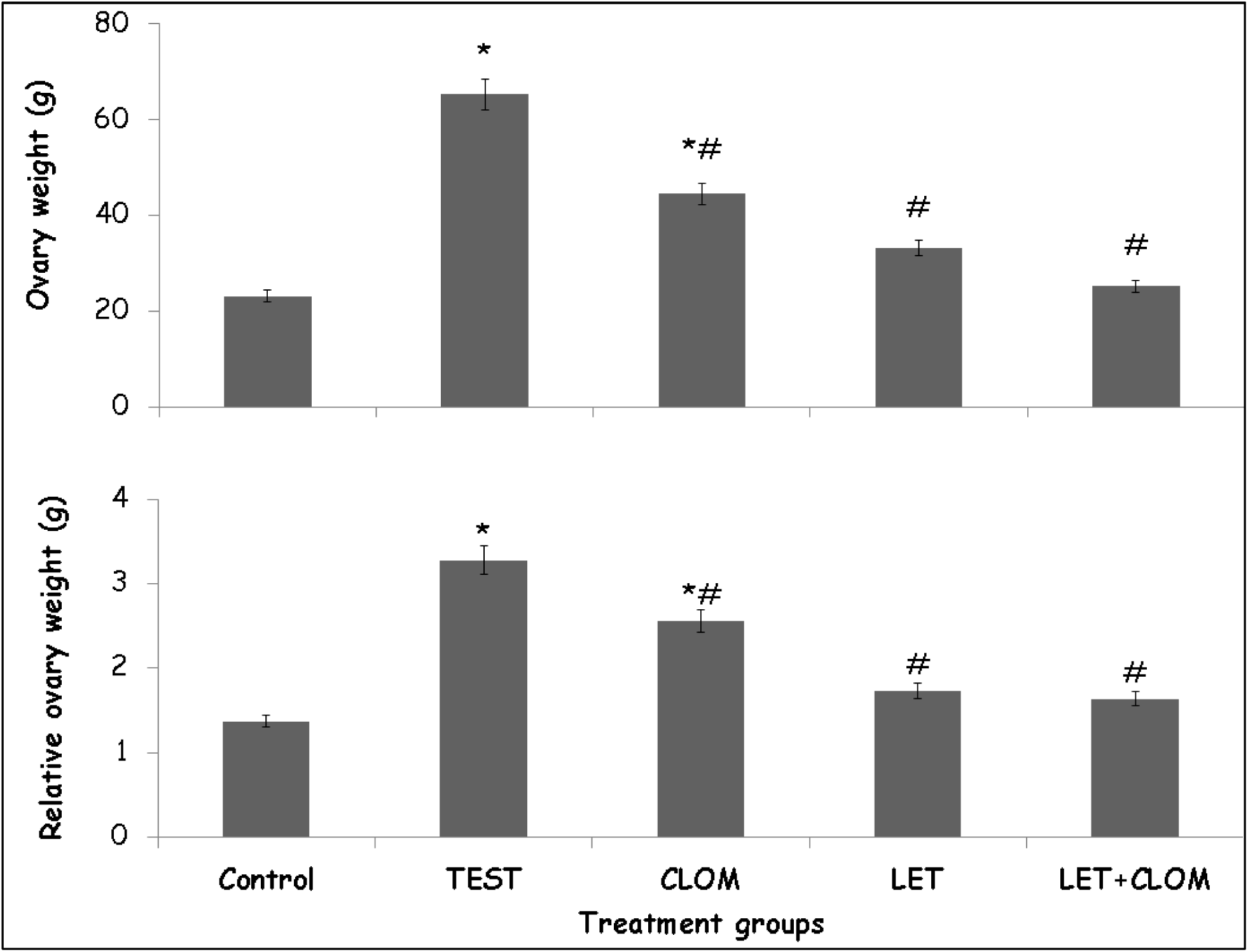
Effect of clomiphene and letrozole on absolute weight of the ovary (upper panel) and relative ovary weight (lower panel) in testosterone treated female rats. Each bar represents Mean ± S.E.M, number of mice per treatment group =6. PCOS: Polycystic ovarian syndrome, CLOM: Clomiphene, LETR: Letrozole.

### 4.5 Effect of clomiphene and letrozole on glucose and lipid levels

Table 1 shows the effect of clomiphene and letrozole on glucose, insulin levels, and lipid parameters in testosterone treated female rats. Glucose levels increased significantly with PCOS, CLOM, LET and CLOM/LET compared to control. Compared to PCOS, glucose levels decreased significantly with CLOM. LET and CLOM/LET.

**Table 1:** Effect of clomiphene letrozole co-administration on glucose and lipid levels.

| Groups | Glucose<br>mg/dl | TC<br>(mmol/L) | TG<br>(mmol/L) | LDL<br>(mmol/L) | HDL (mmol/L) |
| --- | --- | --- | --- | --- | --- |
| Control | 72.4±2.10 | 1.32±0.12 | 0.56±0.10 | 0.58 ±0.21 | 0.32±0.01 |
| TEST | 166.0±2.50* | 7.32±0.35* | 4.14±0.11* | 3.34±0.15* | 1.21±0.02* |
| CLOM | 157.2±1.22*# | 5.10±0.23*# | 3.10±0.1*# | 3.21±0.10*# | 1.11±0.02* |
| LET | 125.2±2.20*# | 3.11±0.20*# | 3.70±0.10*# | 2.61±0.10*# | 0.76±0.03*# |
| CLOM/LET | 88.2±2.13*# | 3.50±0.18*# | 2.18±0.11*# | 2.10±0.11*# | 0.66±0.04*# |

Data represented as Mean ± S.E.M, \*p<0.05 significant difference from control. #p<0.05 significant difference from TEST. number of rats per treatment group =6. TEST: Testosterone, CLOM: Clomiphene, LET: Letrozole.

Total cholesterol levels increased significantly with PCOS, CLOM, LET and CLOM/LET compared with control. Compared to PCOS, total cholesterol levels decreased with CLOM, LET and CLOM/LET. Triglyceride levels increased significantly with PCOS, CLOM, LET and CLOM/LET compared with control. Compared to PCOS, triglyceride levels decreased with CLOM, LET and CLOM/LET.

Low density lipoprotein levels increased significantly with PCOS, CLOM, LET and CLOM/LET compared with control. Compared to PCOS, low density lipoprotein levels decreased with CLOM, LET and CLOM/LET. High density lipoprotein levels increased significantly with PCOS, CLOM, LET and CLOM/LET compared with control. Compared to PCOS, high density lipoprotein levels decreased with CLOM, LET and CLOM/LET.

### 4.5 Effect of clomiphene and letrozole on oestradiol and progesterone levels

Figure 4 shows the effect of clomiphene and letrozole on oestradiol (upper panel) and progesterone (lower panel) levels. Oestradiol levels decreased significantly with PCOS, CLOM and increased with CLOM/LET compared to control. Compared to PCOS, oestradiol levels increased significantly with CLOM, LET and CLOM/LET.

**Figure 4:**
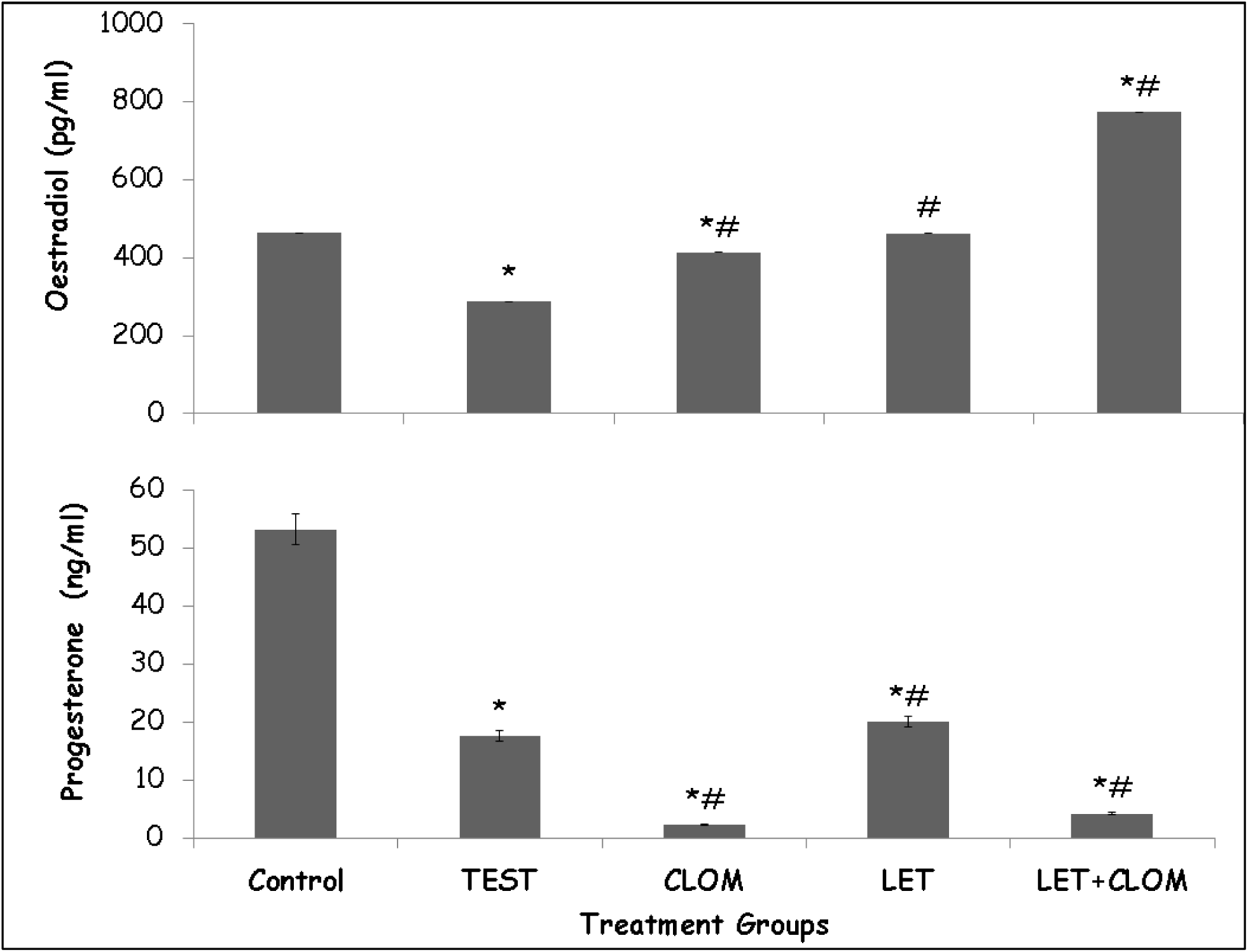
Effect of clomiphene and letrozole on oestrogen (upper panel) and progesterone (lower panel) levels in testosterone treated female rats. Each bar represents Mean ± S.E.M, number of mice per treatment group =6. PCOS: Polycystic ovarian syndrome, CLOM: Clomiphene, LETR: Letrozole.

Progesterone levels decreased significantly with PCOS, CLOM LET and CLOM/LET compared with control. Compared to PCOS, progesterone levels decreased with CLOM, CLOM/LET and increased with LET.

### 4.6 Effect of clomiphene and letrozole on luteinizing hormone and follicle stimulating hormone levels

Figure 5 shows the effect of clomiphene and letrozole on luteinizing hormone (upper panel) and follicle stimulating hormone (lower panel) levels. Luteinizing hormone (LH) levels decreased significantly with PCOS, LET and increased with CLOM/LET compared to control. Compared to PCOS, LH levels increased significantly with CLOM, LET, and CLOM/LET.

**Figure 5:**
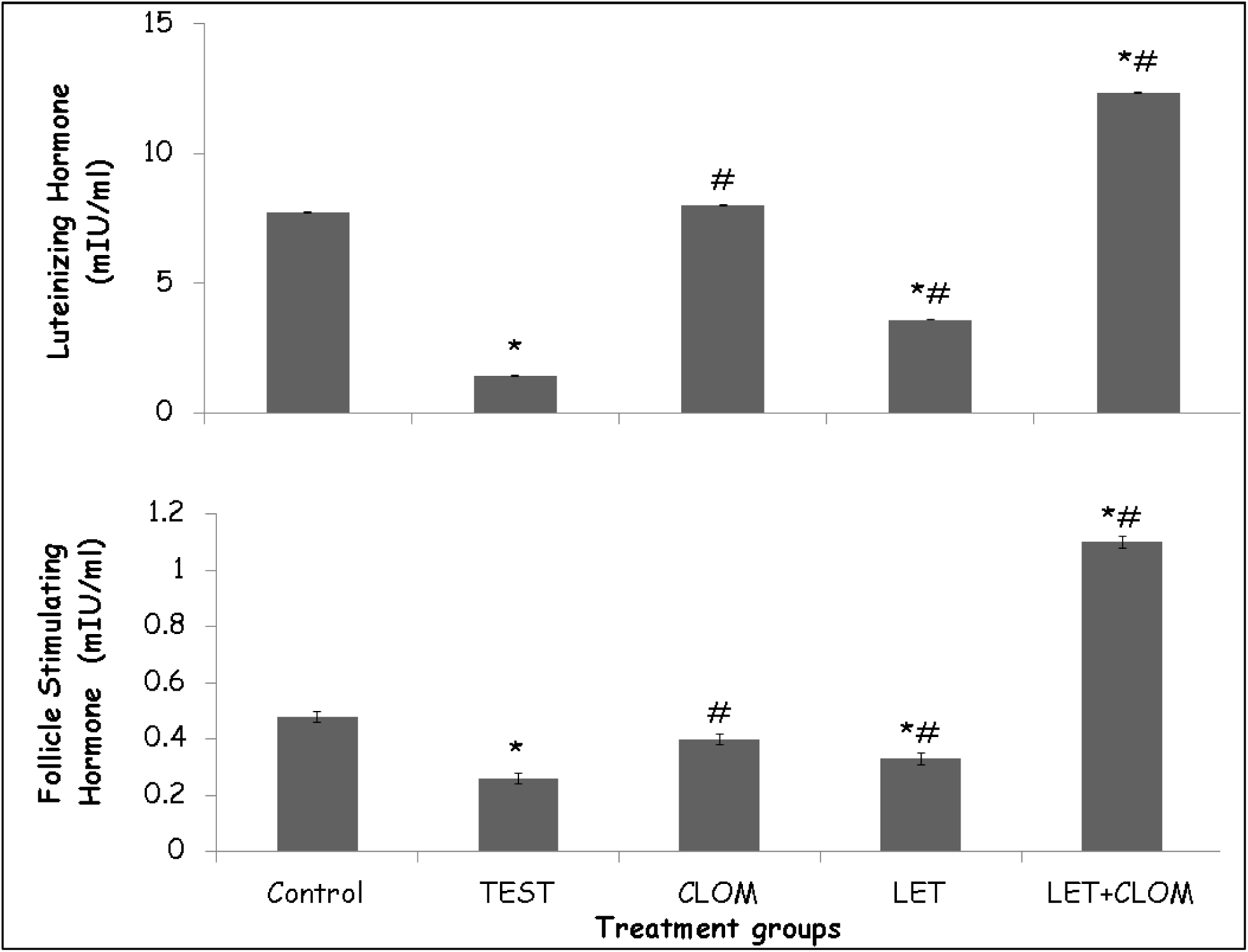
Effect of clomiphene and letrozole on luteinizing (upper panel) and follicle stimulating hormone (lower panel) levels in testosterone treated female rats. Each bar represents Mean ± S.E.M, number of mice per treatment group =6. PCOS: Polycystic ovarian syndrome, CLOM: Clomiphene, LETR: Letrozole.

Follicle stimulating hormone (FSH) levels decreased significantly with PCOS, LET and increased with CLOM/LET compared with control. Compared to PCOS, FSH levels increased with CLOM, LET and CLOM/LET.

### 4.7 Effect of clomiphene and letrozole on lipid peroxidation and Total antioxidant capacity

Figure 6 shows the effect of clomiphene and letrozole on lipid peroxidation levels (upper panel) and total antioxidant capacity (lower panel). Lipid peroxidation levels increased significantly with PCOS, CLOM and CLOM/LET and decreased with LETR compared to control. Compared to PCOS, lipid peroxidation increased significantly with CLOM and decreased with LET and CLOM/LET.

**Figure 6:**
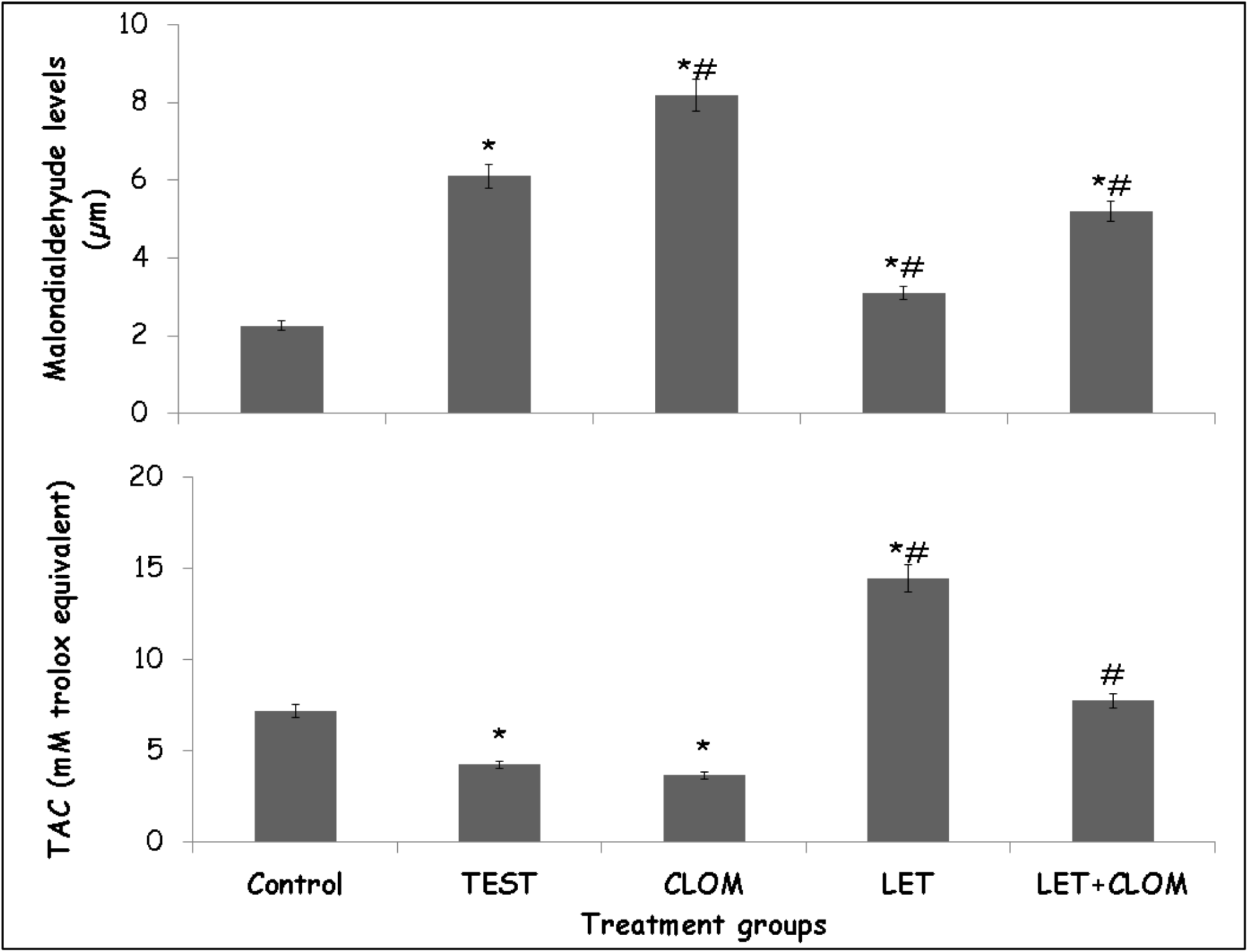
Effect of clomiphene and letrozole on Lipid peroxidation (upper panel) and total antioxidant capacity (lower panel) levels in testosterone treated female rats. Each bar represents Mean ± S.E.M, number of mice per treatment group =6. PCOS: Polycystic ovarian syndrome, CLOM: Clomiphene, LETR: Letrozole.

Total antioxidant capacity decreased significantly with PCOS, CLOM and increased with LET compared with control. Compared to PCOS, total antioxidant capacity increased with LET and CLOM/LET.

### 4.8 Effect of clomiphene and letrozole on interlukin-10 and interleukin 1β levels

Figure 7 shows the effect of clomiphene/ letrozole co administration on interlukin-1β (upper panel) and interleukin 10 (lower panel) levels. Interleukin 1β levels increased significantly with TEST, CLOM and LET compared to control. Compared to TEST, (PCOS control) interleukin 1β levels decreased significantly with CLOM LET and CLOM/LET.

**Figure 7:**
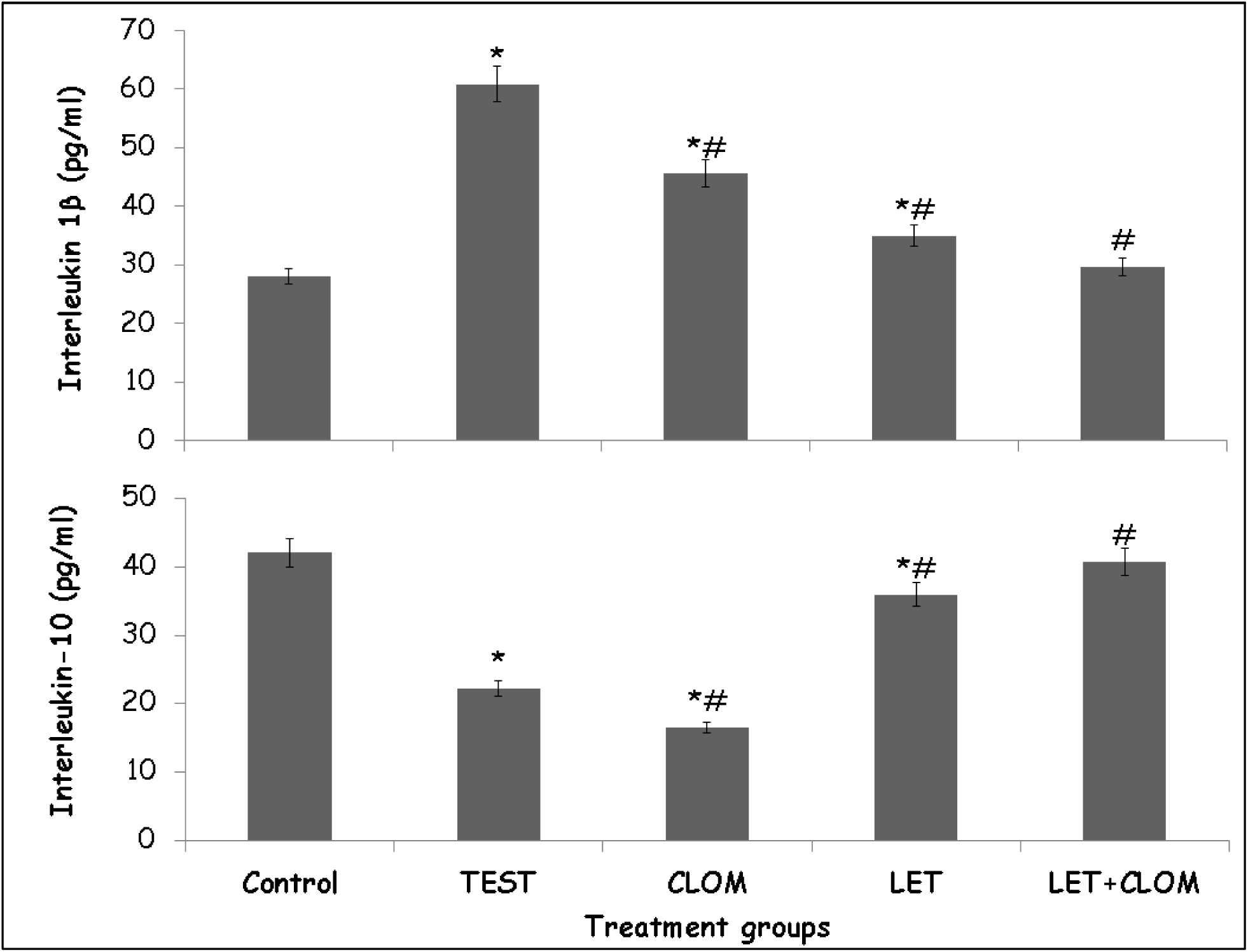
Effect of clomiphene and letrozole on Interleukin-1β (upper panel) and Interleukin-10 (lower panel) levels in testosterone treated female rats. Each bar represents Mean ± S.E.M, number of mice per treatment group =6. PCOS: Polycystic ovarian syndrome, CLOM: Clomiphene, LETR: Letrozole.

Interleukin −10 levels decreased significantly with TEST, CLOM and LET compared with control. Compared to TEST, interleukin 10 levels decreased with CLOM and increased with LET and CLOM/LET.

### 4.9 Effect of clomiphene and letrozole on Tumour necrosis factor-αlevels

Figure 8 shows the effect of clomiphene/letrozole co administration on tumour necrosis factor-α levels. Tumour necrosis factor-α (TNF-α) levels increased significantly with TEST, CLOM, LET and CLOM/LET compared to control. Compared to TEST, (PCOS control), TNF-α levels decreased significantly with CLOM and CLOM/LET.

**Figure 8:**
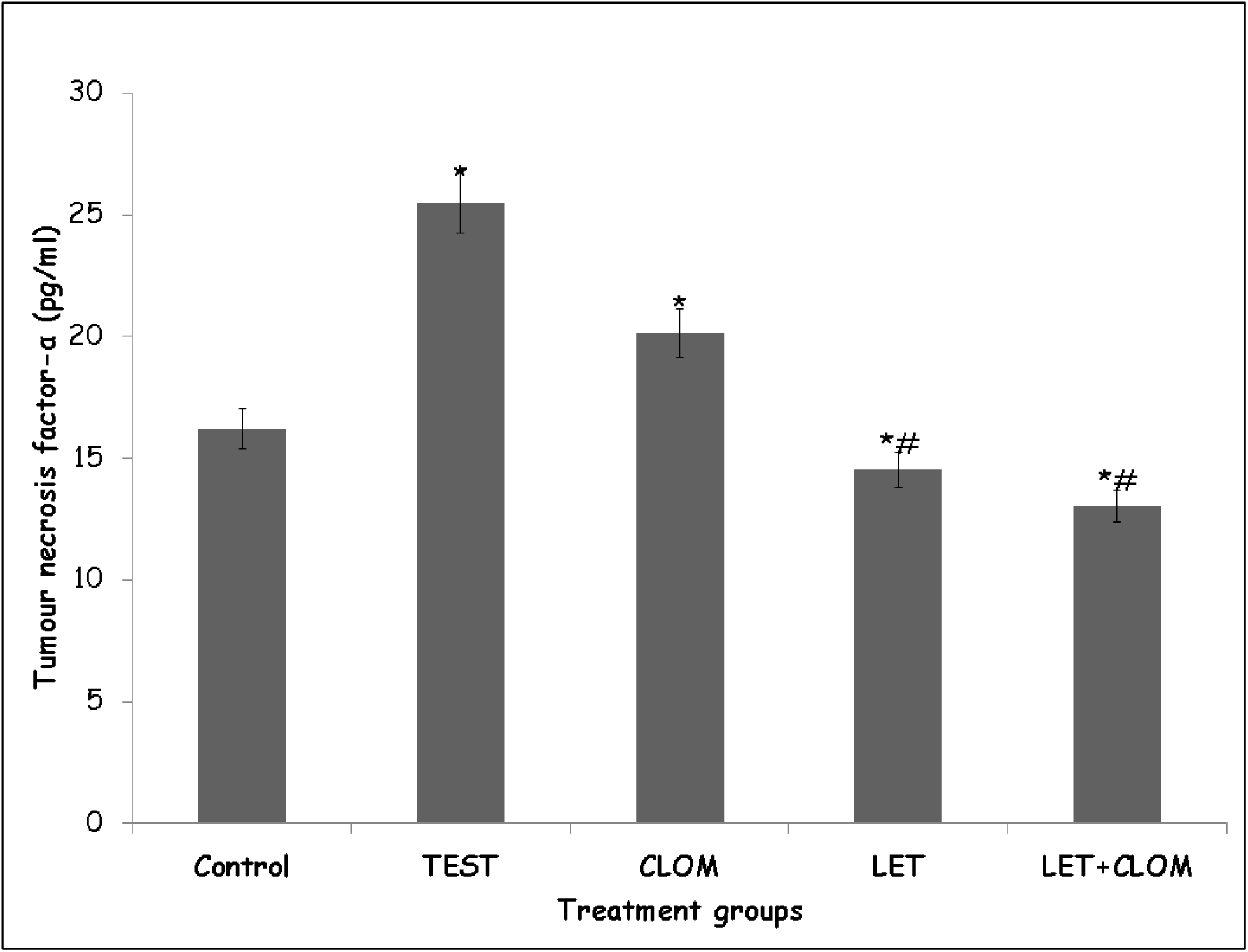
Effect of clomiphene and letrozole on Tumour necrosis factor-α levels in testosterone treated female rats. Each bar represents Mean ± S.E.M, number of mice per treatment group =6. PCOS: Polycystic ovarian syndrome, CLOM: Clomiphene, LETR: Letrozole.

### 4.10 Effect of clomiphene and letrozole on the histology of the ovary (Haematoxylin and Eosin stain)

Figure 9(A-E) shows representative photomicrographs of the haematoxylin and eosin-stained sections of the ovary in testosterone-treated female rats. Ovary of rats in the control group (9A) showed normal follicular architecture with developing antral follicles, intact basement membrane, distinct zona pellucida, and well-formed corpora lutea, in keeping with normal ovarian architecture. In the PCOS control (9B), multiple cystic follicles and thickened basement membrane with numerous secondary/tertiary follicles and reduced corpus luteum was observed, indicating disrupted folliculogenesis. In the clomiphene treatment group (9C), developing follicles (F) with partial restoration of basement membrane integrity and presence of corpora lutea (CL) were observed, suggesting resumption of ovulatory activity. In the letrozole treatment group (9D), cystic and developing follicles with improved follicular structure and active luteinization was observed, while in the combined clomiphene-letrozole treatment group (9E) more organised follicles, compact basement membranes, visible basophilic follicular cells, and multiple corpora lutea, reflecting enhanced restoration of ovarian morphology and follicular development was observed.

**Figure 9:**
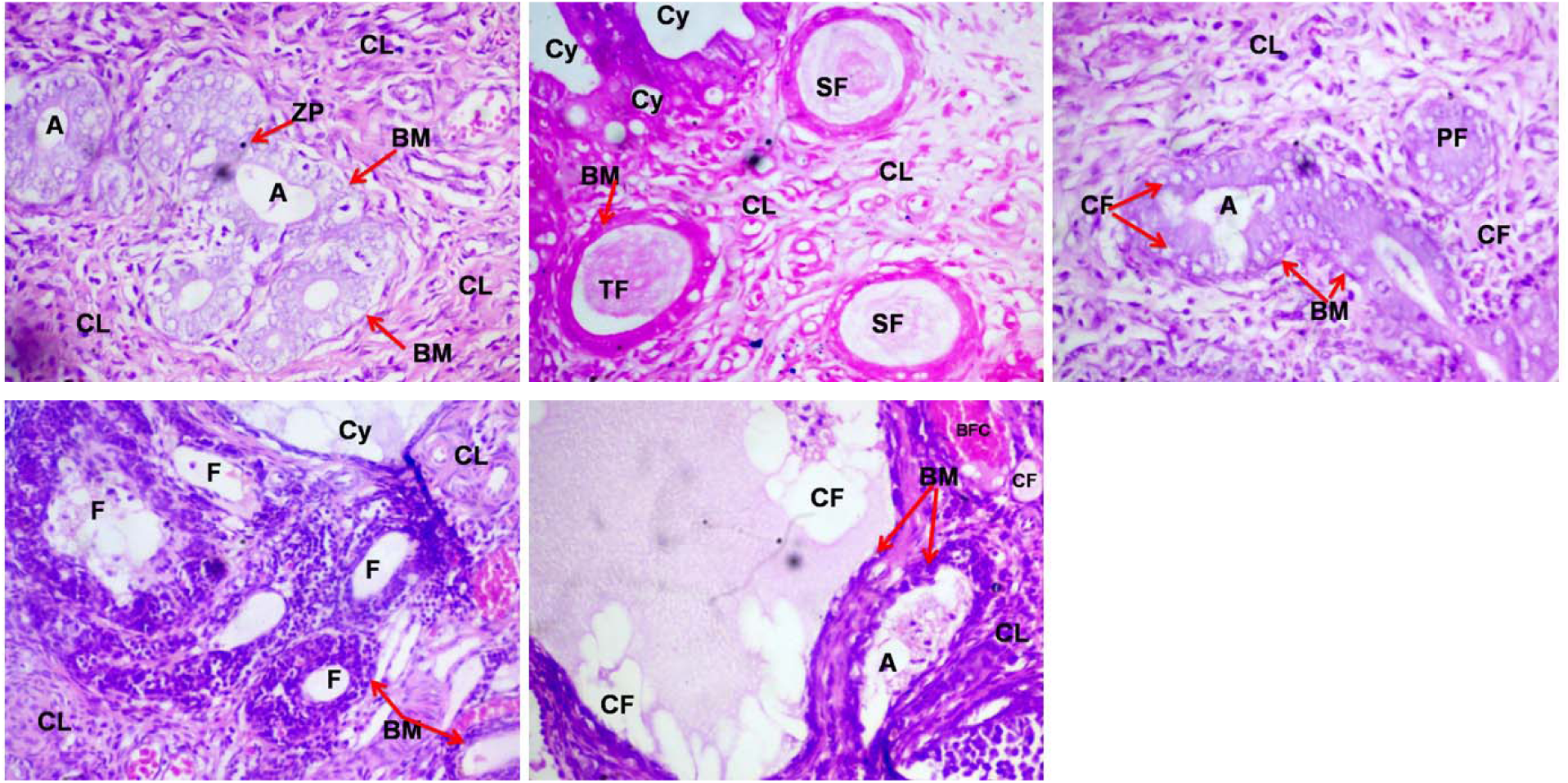
Effect of clomiphene and letrozole on the histology of the ovary (Haematoxylin and Eosin stain) in testosterone treated female rats. Representative photomicrographs of control (A) polycystic ovarian syndrome (PCOS) control (B), PCOS and *clomiphene* (C), PCOS and letrozole (D), PCOS with clomiphene and letrozole (E), A: Antral cells, CL: Corpus luteum, Cy cysts, CF: cystic follicles, F: Follicles, BM: Basement membrane, PF primary follicles, HCF: Haemorrhagic cystic follicles, ZP: Zona pellucida (magnification, 160×)

### 4.11 Effect of clomiphene and letrozole on the histology of the ovary (Haematoxylin and Eosin stain)

Figure 10 demonstrates the ovarian collagen framework visualised with Van Gieson’s stain. In the control group (10A), fine collagen fibres were observed outlining the follicular basement membranes and stroma, representing normal connective tissue distribution. The PCOS control group (10B) displayed intensified red collagen staining, indicating marked stromal fibrosis and thickened follicular basement membranes, a hallmark of chronic ovarian remodeling in PCOS. Following treatment, the clomiphene group (10C) showed mild reduction in collagen deposition, although some areas of fibrosis persisted around atretic follicles. The letrozole group (10D) presented significant improvement, with reduced collagen accumulation and better delineation of follicular architecture. The combined clomiphene and letrozole group (10E) exhibited the least collagen deposition, with collagen distribution similar to that of the control group. The basement membranes appeared thinner and more uniform, reflecting attenuation of fibrosis and restoration of normal stromal structure.

**Figure 10:**
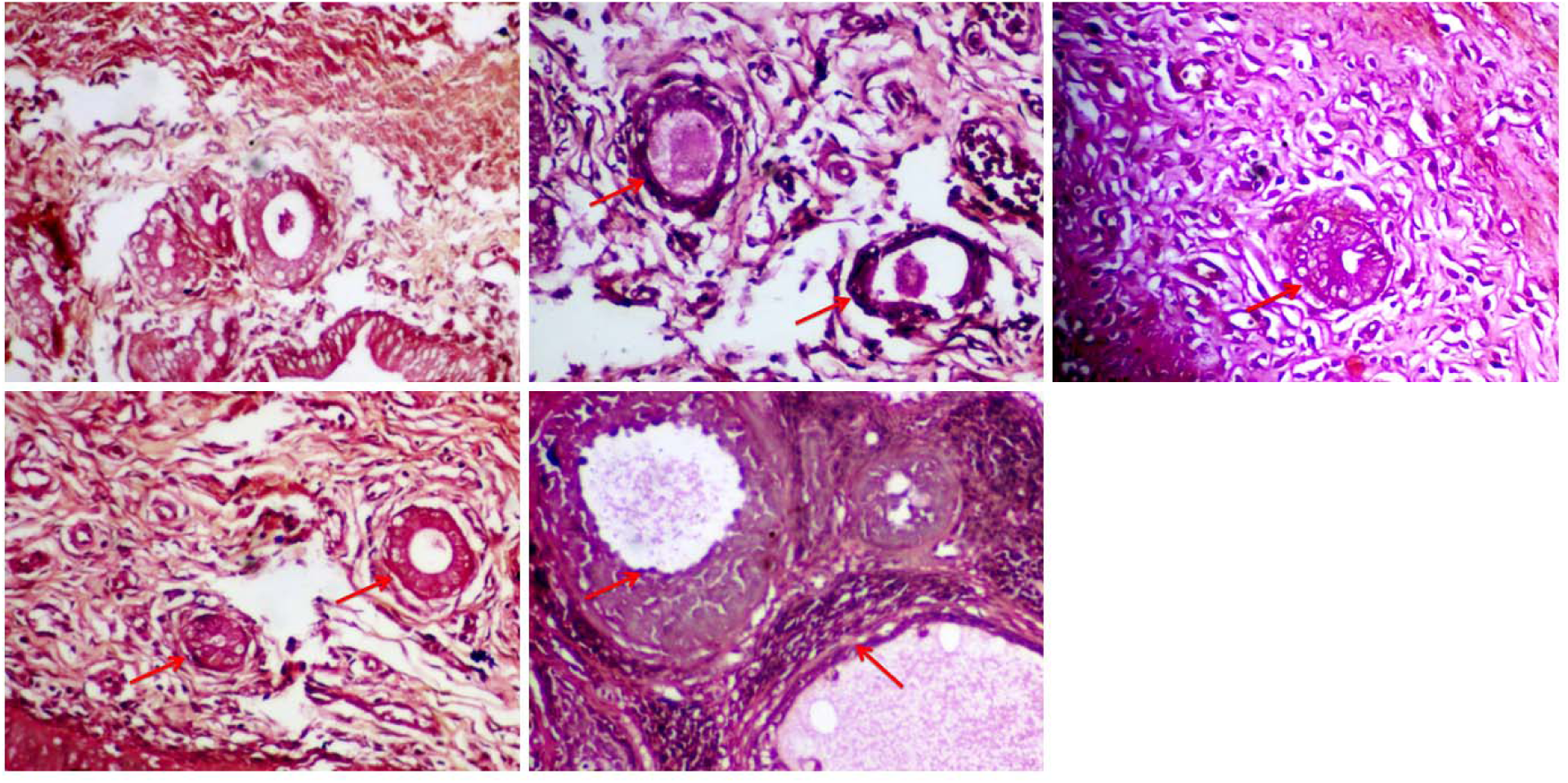
Effect of clomiphene and letrozole on the histology of the ovary (Van Gieson’s stain) in testosterone treated female rats. Representative photomicrographs of control (A) polycystic ovarian syndrome (PCOS) control (B), PCOS and *clomiphene* (C), PCOS and letrozole (D), PCOS with clomiphene and letrozole (E), Red arrows show basement membrane (magnification, 160×)

### 4.12 Effect of clomiphene and letrozole on Ovarian Follicular and Stromal Morphometry

Histomorphometric evaluation using ImageJ software was performed on H&E-stained (Table 2) ovarian sections from testosterone-induced PCOS rats and treatment groups. Quantitative analysis showed that compared to the control group (9A), rats in the PCOS control group (9B) exhibited marked reduction in the number of developing follicles, a decrease in mean follicular diameter, thickened basement membranes, and complete absence of corpora lutea, confirming follicular arrest and anovulation. Rats in the clomiphene group (9C) showed in a modest increase in follicular count and follicular diameter, with partial restoration of antral follicles when compared with control, but a decrease when compared with PCOS control, while the letrozole treatment (9D) produced revealed a significant increase in mean follicular diameter and reappearance of growing follicles compared to PCOS control. The combination of clomiphene and letrozole (9E) produced the most pronounced histological recovery, with multiple active follicles, well-formed corpora lutea, and a normalised basement membrane thickness comparable to control group.

**Table 2:** Effect of clomiphene and letrozole on Ovarian Follicular and Stromal Morphometry (H& E)

| Group | Follicular count (per field) | Mean follicular diameter ($\mu\text{m}$) | Basement membrane thickness ($\mu\text{m}$) |
| --- | --- | --- | --- |
| Control | $8.6 \pm 0.4$ | $128.3 \pm 7.2$ | $1.8 \pm 0.3$ |
| TEST | $3.2 \pm 0.6^*$ | $78.4 \pm 5.1^*$ | $4.7 \pm 0.5^*$ |
| CLOM | $5.9 \pm 0.7^{* \#}$ | $103.5 \pm 6.8^{* \#}$ | $3.2 \pm 0.4^{* \#}$ |
| LET | $7.1 \pm 0.5^{* \#}$ | $115.2 \pm 8.3^{* \#}$ | $2.6 \pm 0.3^{* \#}$ |
| CLOM/LET | $8.4 \pm 0.6^{\#}$ | $126.7 \pm 7.1^{\#}$ | $1.9 \pm 0.2^{\#}$ |

Data represented as Mean ± S.E.M, \* $p < 0.05$ significant difference from control. # $p < 0.05$ significant difference from TEST. number of rats per treatment group =6. TEST: Testosterone, CLOM: Clomiphene, LET: Letrozole.

**Table 3:** Effect of Clomiphene and Letrozole on Collagen Fibre Density (Van Gieson Sections)

| Group | Collagen area (%) | Collagen fibre intensity (a.u.) |
| --- | --- | --- |
| Control | $21.4 \pm 2.3$ | $42.8 \pm 4.1$ |
| TEST | $49.7 \pm 3.8^*$ | $91.2 \pm 5.3^*$ |
| CLOM | $34.5 \pm 2.9^{* \#}$ | $63.7 \pm 4.2^{* \#}$ |
| LET | $29.8 \pm 3.2^{* \#}$ | $55.1 \pm 3.7^{* \#}$ |
| CLOM/LET | $23.6 \pm 2.5^{\#}$ | $44.9 \pm 3.9^{\#}$ |

Data represented as Mean ± S.E.M, \* $p < 0.05$ significant difference from control. # $p < 0.05$ significant difference from TEST. number of rats per treatment group =6. TEST: Testosterone, CLOM: Clomiphene, LET: Letrozole. A.u : arbitrary units

### 4.13 Effect of clomiphene and letrozole on Collagen Fiber Density (Van Gieson Sections)

Von Gieson staining revealed distinct alterations in stromal collagen architecture among groups. PCOS control ovaries displayed marked perivascular and perifollicular collagen deposition with irregular thickening of the basement membrane, indicating fibrotic remodeling. Quantitative ImageJ colour thresholding (red-channel segmentation) showed that collagen area percentage was significantly elevated in the PCOS group compared to control. Treatment with clomiphene or letrozole individually reduced collagen deposition, while the combination therapy produced near-normal collagen patterns similar to controls.

## Discussion

Polycystic ovary syndrome (PCOS) is a complex metabolic and hormonal disorder characterised by features such as obesity, hirsutism, acne, irregular menstrual cycles, and insulin resistance. In this study, the results revealed that the co-administration of clomiphene citrate and letrozole significantly reduced body and ovary weights, lowered glucose levels, and improved oestradiol and follicle-stimulating hormone profiles compared with PCOS control. The combination therapy also enhanced antioxidant capacity, reduced lipid peroxidation, and modulated inflammatory cytokines by lowering IL-1β and TNF-α while elevating IL-10. Histological evaluation revealed cystic follicles with basement membrane thickening in the PCOS group rats, consistent with ovarian hyperstimulation; and variable levels of reversal of these features with clomiphene and/or letrozole treatment.

Increased body weight in PCOS has been described as both a cause and an effect, with obesity known to contribute both to the development and also the worsening of PCOS; especially in relation to features such as insulin resistance and high androgen levels. Also, studies have reported that the hormonal and metabolic imbalances that characterise PCOS make weight loss difficult, resulting a cyclical relationship [31–34]. In this study, testosterone administration induced a significant increase in body weight in female rats, consistent with previous reports linking hyperandrogenism with increased adiposity and metabolic dysfunction in PCOS models [35, 36]. These features have also been attributed to androgen’s influence on energy balance, feed intake and body composition, resulting in increased fat accumulation [31–34]. The marked weight gain in the PCOS control group highlights the metabolic consequences of androgen excess, which include central obesity and insulin resistance, both hallmarks of PCOS pathophysiology [37]. Interestingly, while all treatment groups (TEST, CLOM, LETR, CLOM/LETR) showed weight gain relative to normal controls, animals treated with clomiphene and letrozole exhibited a slower trajectory of weight increase. This trend suggests a partial amelioration of androgen-induced metabolic alterations. Notably, weight gain was significantly elevated in the PCOS and LETR groups, whereas it was reduced in the CLOM and CLOM/LETR groups. The combination therapy (CLOM/LETR) demonstrated the most pronounced reduction in relative weight gain when compared with the PCOS group, suggesting a potential additive or synergistic effects of the two agents in modulating body weight. These observations align with earlier studies reporting that clomiphene and aromatase inhibitors may indirectly influence metabolic parameters through modulation of estrogen-androgen balance, improvement of gonadotrophin secretion, and enhancement of ovarian function [38, 39]. The greater effect of the CLOM/LETR combination implies that combined targeting of hypothalamic–pituitary oestrogen signaling and peripheral oestrogen synthesis may attenuate the metabolic dysregulation associated with hyperandrogenism more effectively than monotherapy. Overall, the results indicate that while testosterone induces significant weight gain in female rats, clomiphene, especially in combination with letrozole can mitigate this effect, pointing to possible advantages of dual therapy in managing not only reproductive but also metabolic aspects of PCOS

The relationship between feed intake and PCOS is complex. In this study, the effects of clomiphene and letrozole on feed intake highlight important differences in how these agents influence feeding behaviour in testosterone-treated female rats. Weekly feed intake was significantly elevated in the CLOM and LETR groups compared with both the normal and PCOS controls, while co-administration of CLOM/LETR produced a significant reduction in feed intake across the study period. Rats in the combination group consistently consumed the least feed, indicating a potential suppressive effect on appetite or energy intake when the two agents are used together. The finding supports studies that have reported increased feed intake with clomiphene therapy [40]. Analysis of relative changes in feed intake further reinforced this trend. Percentage change increased significantly in the CLOM and LETR groups compared to controls, but was significantly reduced in the CLOM/LETR group when compared to both normal and PCOS controls. These findings suggest that clomiphene and letrozole, when administered individually, may enhance feeding behaviour, possibly due to their modulatory effects on oestrogen and gonadotrophin signaling, which are known to influence appetite and metabolic regulation. However, the combination treatment appears to counteract this effect, normalising or even suppressing feed intake relative to the PCOS group. The reduced feed intake in the CLOM/LETR group could explain the attenuated body weight gain observed, pointing to an interplay between appetite regulation and metabolic outcomes under dual therapy. This finding is particularly relevant given that hyperphagia and weight gain exacerbate the metabolic and reproductive complications of PCOS [41, 42]. Thus, the combination of clomiphene and letrozole may offer advantages not only in reproductive endocrinology but also in modulating energy balance.

In PCOS, in addition to the ovaries being populated with underdeveloped ovarian follicles creating the "polycystic" appearance, there have also been accounts of them being enlarged [43]. In this study, ovary weight and relative ovary weight increased significantly in the PCOS control group compared to normal control, reflecting ovarian enlargement commonly associated with hyperandrogenism. Interestingly, clomiphene monotherapy also resulted in a significant increase in ovarian weight compared with control, which may be attributed to its stimulatory effects on gonadotropin release and follicular recruitment. When compared with the PCOS group, however, ovary weight decreased significantly in all treatment groups (CLOM, LETR, CLOM/LETR), suggesting that both monotherapies and the combination therapy exerted a normalising influence on ovarian hypertrophy. A similar pattern was observed for relative ovary weight, which was significantly increased in PCOS and CLOM groups compared with control, but reduced in all other treated groups relative to PCOS. These findings indicate that while clomiphene alone may transiently increase ovarian mass due to follicular stimulation, its long-term effect, especially when combined with letrozole, tends to counteract testosterone-induced ovarian enlargement [43]. Letrozole, by reducing oestrogen synthesis and modulating androgen excess, may contribute to this protective effect. The reduction in ovary weight with the CLOM/LETR combination suggests a synergistic action, potentially reflecting improved regulation of folliculogenesis and attenuation of cystic ovarian changes.

Polycystic ovary syndrome has been associated with impaired glucose tolerance and dysmetabolism [44–46]. In this study, hyperandrogenism induced by testosterone significantly elevated glucose, total cholesterol, triglyceride, LDL, and HDL levels compared with controls, confirming the development of metabolic disturbances consistent with PCOS. These findings mirror clinical reports where PCOS is strongly associated with impaired glucose tolerance, dyslipidaemia and increased cardiometabolic risk [45, 46]. Treatment with clomiphene, letrozole, or their combination significantly reduced glucose levels compared to the PCOS group, with the most pronounced effect observed in the combination therapy. This suggests that both agents may contribute to improved insulin sensitivity and glucose regulation, possibly through modulation of ovarian steroidogenesis and gonadotrophin secretion. Similarly, all lipid parameters (total cholesterol, triglycerides, LDL, and HDL) were significantly elevated in the PCOS group compared with controls, consistent with androgen-induced dyslipidemia [47]. However, treatment with clomiphene, letrozole, or their combination resulted in significant reductions across these lipid measures compared with PCOS. This pattern indicates that ovulation-inducing agents may exert secondary benefits on lipid metabolism, likely mediated through improvements in insulin sensitivity, oestrogen-androgen balance, and reduced oxidative stress. Although HDL (traditionally considered cardioprotective) was elevated in PCOS rats, its subsequent reduction following treatment may reflect normalisation of lipid turnover rather than a detrimental outcome. Taken together, these results suggest that clomiphene and letrozole, particularly when used in combination, not only improve reproductive endpoints but also confer beneficial effects on glucose and lipid metabolism in PCOS.

Result of hormonal assays revealed hormonal alterations, alongside derangements of inflammatory cytokine and oxidative stress profiles. These provide important mechanistic insights into the effects of clomiphene and letrozole in hyperandrogenic rats. Reduced oestradiol and progesterone in the PCOS group reflect disrupted folliculogenesis, anovulation, and luteal insufficiency, consistent with clinical PCOS [48, 49]. Restoration of oestradiol with CLOM, LETR, and particularly CLOM/LETR, suggests partial recovery of follicular activity. However, persistently reduced progesterone, especially in the CLOM and CLOM/LETR groups, implies that follicular recruitment did not consistently translate into effective luteinization, a phenomenon supported by the histological evidence of multiple cystic follicles with thickened basement membranes. These endocrine disruptions were accompanied by significant changes in oxidative stress markers. PCOS rats showed elevated lipid peroxidation and reduced total antioxidant capacity, indicating a pro-oxidant state consistent with androgen-induced mitochondrial dysfunction [50, 51]. Treatment with LETR and CLOM/LETR significantly reduced lipid peroxidation and increased antioxidant capacity, suggesting that letrozole, either alone or in combination, is more effective at restoring redox balance than clomiphene alone. Improved oestradiol levels in these groups may also contribute, as oestrogens possess intrinsic antioxidant properties [52]. The immune-inflammatory axis further reinforced this pattern; as PCOS was associated with elevated pro-inflammatory cytokines (IL-1β, TNF-α) and reduced anti-inflammatory IL-10, consistent with chronic low-grade inflammation characteristic of PCOS [53] However, CLOM/LETR treatment significantly suppressed IL-1β and TNF-α while increasing IL-10, indicating an anti-inflammatory effect beyond that seen with either monotherapy. These changes may be linked to restored oestrogen signaling and improved oxidative balance, as both oestradiol and antioxidant pathways are known to suppress inflammatory cascades [54]. Taken together, the combination of clomiphene and letrozole appears to modulate the endocrine–oxidative–immune axis more effectively than either drug alone. While dual therapy enhances follicular activity (increased oestradiol) and mitigates oxidative stress and inflammation, its relative suppression of progesterone suggests that ovarian hyperstimulation may come at the cost of luteal function [55]. This highlights both the therapeutic promise and potential limitations of combined ovulation induction strategies in PCOS.

In PCOS, the hypothalamic-pituitary-ovarian axis is disrupted, leading to abnormal gonadotropin levels as observed in this study. In the PCOS groups, both LH and FSH levels were significantly reduced compared with controls, indicating disruption of the hypothalamic–pituitary-ovarian axis. While this finding differs slightly from the classic clinical PCOS phenotype, where LH is often elevated relative to FSH, it aligns with the suppressed gonadotrophin release reported in androgen-driven rodent models, due to negative feedback inhibition [56, 57]. Treatment with clomiphene or letrozole restored LH and FSH levels significantly compared with the PCOS group, consistent with their mechanisms of action. Clomiphene, through oestrogen receptor antagonism at the hypothalamus, enhances GnRH pulsatility and gonadotrophin release [57]. Letrozole, via aromatase inhibition, reduces circulating estrogens, thereby relieving negative feedback and increasing FSH secretion. Notably, combination therapy (CLOM/LETR) produced the most pronounced increases in both LH and FSH relative to PCOS, indicating a possible synergistic effect on pituitary stimulation. The rise in FSH is particularly important for follicular recruitment and maturation, while increased LH supports theca cell androgen production and subsequent aromatization to estrogens. This hormonal milieu explains the observed increase in estradiol with CLOM/LETR, as multiple follicles were likely recruited. However, the persistently low progesterone in CLOM and CLOM/LETR groups suggests inadequate luteinization despite improved gonadotrophin drive. These findings reinforce that clomiphene and letrozole act at different levels of the HPG axis, and their combination enhances gonadotrophin release more effectively than monotherapy [58, 59]. This synergism supports follicular development and oestradiol synthesis, but the imbalance between estrogen and progesterone may predispose to ovarian hyperstimulation, as seen in the histological profiles.

The hormonal profile of testosterone-induced PCOS rats revealed profound disruption of the hypothalamic–pituitary–ovarian (HPO) axis. In the PCOS group, oestradiol and progesterone levels were significantly reduced compared with controls, indicating impaired folliculogenesis, anovulation, and luteal dysfunction. Similarly, both LH and FSH levels decreased in PCOS animals, reflecting suppression of gonadotrophin release, likely mediated by negative feedback from excess androgens. Although this contrasts with the elevated LH/FSH ratio often reported in clinical PCOS, it aligns with androgen-driven rodent models, where chronic hyperandrogenism dampens hypothalamic–pituitary responsiveness. Treatment with clomiphene and letrozole modulated these endocrine derangements in complementary ways. Clomiphene, through oestrogen receptor antagonism, enhanced gonadotrophin secretion and partially restored estradiol. Letrozole, via aromatase inhibition, reduced oestrogenic negative feedback, increasing FSH release and follicular recruitment [60]. Combination therapy (CLOM/LETR) produced the most marked increases in both LH and FSH, suggesting a synergistic stimulatory effect on pituitary output. This robust gonadotrophin drive was accompanied by a significant rise in oestradiol, particularly in the CLOM/LETR group, consistent with enhanced follicular growth and recruitment. However, progesterone levels remained markedly suppressed across all groups, with the lowest levels observed in CLOM and CLOM/LETR-treated animals. This suggests that while follicular development was improved, luteinization and corpus luteum formation were inadequate, likely reflecting the recruitment of multiple immature follicles rather than successful ovulation and luteal transformation. This hormonal imbalance characterized by increased oestradiol but persistently low progesterone may predispose to ovarian hyperstimulation, a finding supported by the ovarian histomorphology. Overall, these results demonstrate that clomiphene and letrozole, particularly in combination, effectively restore gonadotrophin secretion and oestradiol synthesis but do not adequately normalise progesterone, highlighting the complexity of re-establishing full ovarian endocrine competence in PCOS

Also, in this study, increased oxidative stress was observed with testosterone-induced PCOS rats with a modulation of these effects with clomiphene and letrozole. Lipid peroxidation (measured as malondialdehyde (MDA) levels in this study) which is an indicator of oxidative damage to cell membranes, was significantly elevated in the PCOS group compared with control, reflecting the heightened pro-oxidant state associated with hyperandrogenism and metabolic imbalance. A similar elevation in MDA was observed with clomiphene treatment, while CLOM/LETR treatment showed a reduction relative to PCOS but remained higher than controls. In contrast, letrozole monotherapy markedly reduced lipid peroxidation compared with both control and PCOS, suggesting a stronger antioxidant protective effect. Total antioxidant capacity (TAC) followed an inverse trend. PCOS animals exhibited reduced TAC, consistent with diminished endogenous antioxidant defense. Clomiphene also suppressed TAC, indicating limited ability to restore redox balance. By contrast, letrozole treatment significantly enhanced TAC relative to both PCOS and controls, while CLOM/LETR improved TAC compared with PCOS but to a lesser extent than letrozole alone. These findings suggest that letrozole is more effective than clomiphene in correcting oxidative imbalance in PCOS. Letrozole’s ability to reduce lipid peroxidation and boost antioxidant reserves may be attributed to its suppression of oestrogen-driven reactive oxygen species generation and improved metabolic profile [61]. The combination therapy, while partially beneficial, appears less protective than letrozole alone, possibly reflecting opposing interactions between clomiphene’s pro-oxidant effects and letrozole’s antioxidant actions. Taken together, these results reinforce the concept that oxidative stress is a central mediator of PCOS pathology and highlight differential drug effects: clomiphene primarily modulates gonadotrophin and follicular dynamics but fails to correct redox imbalance, whereas letrozole exerts both endocrine and antioxidant benefits.

Alterations in the pro- and anti-inflammatory cytokine were observed in the groups with testosterone-induced PCOS, while administration of letrozole and/or clomiphene modulated these effects. Interleukin-1β (IL-1β), a key pro-inflammatory cytokine implicated in ovarian dysfunction and metabolic disturbances in PCOS [63, 64], was significantly elevated in the PCOS group compared with controls, confirming a heightened inflammatory milieu. Both clomiphene and letrozole monotherapies also showed elevated IL-1β compared with controls; however, relative to PCOS, IL-1β levels were significantly reduced with each treatment, and further suppressed when the drugs were co-administered. This suggests that while both agents attenuate hyperandrogenism-driven inflammation, combination therapy provides a more robust downregulation of IL-1β. In contrast, interleukin-10 (IL-10), an anti-inflammatory cytokine, was significantly reduced in PCOS, clomiphene, and letrozole groups compared with control, reflecting compromised immunoregulatory capacity. Importantly, compared to PCOS, IL-10 levels increased significantly with letrozole and CLOM/LETR co-treatment, whereas clomiphene alone failed to improve IL-10. These findings highlight letrozole’s stronger immunomodulatory potential, consistent with its antioxidant profile. The partial restoration of IL-10 with CLOM/LETR suggests that letrozole’s anti-inflammatory actions can counterbalance clomiphene’s limited effect. Taken together, the cytokine results indicate that PCOS is characterized by a pro-inflammatory shift (↑IL-1β, ↓IL-10), which is incompletely corrected by clomiphene but more effectively modulated by letrozole and the CLOM/LETR combination. These findings align with the oxidative stress results, where letrozole exerted superior antioxidant and cytoprotective effects. Collectively, they reinforce the emerging view of PCOS as a state of oxidative–inflammatory imbalance and suggest that letrozole, alone or in combination with clomiphene, may confer added benefits by targeting both endocrine and immune dysregulation.

Histological analysis of ovarian sections further clarifies the functional impact of testosterone-induced PCOS and its modulation by clomiphene, letrozole, and their combination. Control ovaries displayed normal architecture, with clearly identifiable antral follicles, oocytes, zona pellucida, corpus luteum, and intact basement membranes, features consistent with normal ovulatory cycles. In contrast, PCOS ovaries showed classic pathological changes: absence of corpus luteum (indicative of anovulation), multiple cystic follicles, excessive primary follicles, and thickened basement membranes. These alterations mirror the endocrine derangements observed (reduced oestradiol, progesterone, and gonadotrophin secretion) and confirm disrupted folliculogenesis and ovulation. Treatment with clomiphene or letrozole partially restored ovarian morphology. Both groups demonstrated the presence of corpus luteum and structurally organised basement membranes, indicating some degree of follicular maturation and ovulation. However, a few cystic follicles persisted, reflecting incomplete reversal of the PCOS phenotype. Notably, co-administration of clomiphene and letrozole resulted in exaggerated ovarian changes. Although corpus luteum and antral cells were present, there were also numerous small to very large cystic follicles and pronounced basement membrane thickening, consistent with ovarian hyperstimulation. This aligns with the endocrine data, where CLOM/LETR markedly elevated gonadotrophins and oestradiol but failed to normalise progesterone. The persistence of immature or cystic follicles despite enhanced hormonal drive suggests dysregulated follicular recruitment without adequate luteinization. Collectively, these histological findings highlight that while clomiphene and letrozole can independently ameliorate ovarian dysfunction in PCOS, their combined use may induce hyperstimulation syndrome–like pathology, raising concerns about their translational safety and reinforcing the need for cautious dosing and monitoring.

Histomorphometric evaluation of ovarian sections revealed clear pathological alterations in testosterone-induced PCOS rats, consistent with features of disrupted folliculogenesis and stromal fibrosis. The PCOS control group showed a marked reduction in developing follicles, decreased follicular diameter, and thickened basement membranes, alongside the absence of corpora lutea an indication of arrested follicular maturation and anovulation. These observations align with the established pathophysiology of PCOS, where hyperandrogenism and altered gonadotropin dynamics disrupt follicular recruitment and selection, leading to accumulation of immature follicles and loss of cyclic ovulation. Treatment with clomiphene citrate induced partial histological recovery, evidenced by a moderate increase in follicular count and diameter, as well as partial reappearance of antral follicles. Clomiphene’s anti-estrogenic mechanism at the hypothalamic level enhances gonadotropin release, particularly FSH, thereby promoting follicular growth. However, its incomplete restoration of follicular structure in this study suggests potential limitations in reversing testosterone-induced stromal remodeling, possibly due to its prolonged anti-oestrogenic effects on ovarian tissue. In contrast, letrozole-treated ovaries demonstrated a more-pronounced improvement in follicular morphology, with significant increases in mean follicular diameter and reappearance of growing follicles. As a selective aromatase inhibitor, letrozole reduces peripheral oestrogen production, lowering negative feedback on the hypothalamic–pituitary axis and stimulating endogenous FSH secretion [65]. This mechanism likely underlies the enhanced follicular recruitment observed, consistent with clinical and preclinical reports suggesting superior ovulatory outcomes with letrozole compared to clomiphene. The combination therapy (CLOM/LETR) produced the most-remarkable histological recovery, featuring multiple developing follicles, visible corpora lutea, and normalised basement membrane thickness comparable to the control group. This synergistic effect suggests complementary mechanisms where letrozole optimises FSH stimulation while clomiphene extends the follicular response window resulting in improved folliculogenesis and ovulatory restoration.

Complementary findings from Van Gieson–stained sections further underscore the therapeutic impact of the treatments on ovarian stromal integrity. The PCOS control group exhibited dense perivascular and perifollicular collagen deposition, reflecting stromal fibrosis and altered extracellular matrix (ECM) remodeling, hallmark features of chronic hyperandrogenic states. Quantitative ImageJ analysis confirmed a significant elevation in collagen area percentage in PCOS rats relative to controls. Both clomiphene and letrozole monotherapies mitigated collagen accumulation, while combination therapy effectively normalised collagen distribution and reduced stromal fibrosis to near-control levels. This suggests that dual modulation of oestrogenic and aromatase pathways not only restores follicular morphology but also reverses fibrotic remodeling, enhancing overall ovarian microarchitecture. Taken together, these histomorphometric and collagen analyses demonstrate that combined clomiphene and letrozole therapy exerts a superior corrective effect on the ovarian structure of PCOS rats compared with either agent alone. The observed recovery in follicular development and reduction in stromal fibrosis suggest a restoration of normal cyclic ovarian function. These findings provide strong preclinical evidence supporting the potential benefit of combination ovulation-induction therapy in PCOS, particularly in cases resistant to monotherapy.

## Conclusion

In summary, this study provides compelling experimental evidence that combined administration of clomiphene citrate and letrozole has superior therapeutic efficacy compared to monotherapy in a testosterone-induced rat model of PCOS. The combination therapy effectively mitigated hyperandrogenism-associated metabolic and endocrine disturbances by improving oestradiol and follicle-stimulating hormone levels, reducing body and ovarian weights, and normalisng glucose and lipid profiles. Furthermore, it enhanced total antioxidant capacity, attenuated lipid peroxidation, and modulated inflammatory cytokines decreasing IL-1β and TNF-α while restoring anti-inflammatory IL-10.

Histological and histomorphometric analyses corroborated these biochemical findings, revealing restoration of follicular development, reappearance of corpora lutea, and normalisation of basement membrane thickness. Van Gieson staining further demonstrated reduced stromal collagen deposition, indicating reversal of ovarian fibrosis and improved microarchitectural integrity. Collectively, these outcomes suggest that dual modulation of hypothalamic–pituitary– gonadal feedback and peripheral aromatase activity underlies the combined effect of clomiphene and letrozole on ovarian function.

Despite these benefits, the persistence of reduced progesterone levels and evidence of mild ovarian hyperstimulation in the combination group warrant careful consideration in potential clinical translation. Nonetheless, this dual-therapy approach holds promise as a more-comprehensive strategy for managing both the reproductive and metabolic dimensions of PCOS. Future studies should explore optimal dosing strategies, long-term safety profiles, and translational validation in clinical settings, particularly among patients with clomiphene-resistant or metabolically-severe forms of PCOS.

### Translational Relevance Statement

This study highlights the therapeutic potential of co-administering clomiphene citrate and letrozole in managing both the reproductive and metabolic abnormalities associated with polycystic ovary syndrome (PCOS). By demonstrating possible synergistic improvements in hormonal balance, oxidative/inflammatory regulation, and ovarian histoarchitecture; this work provides mechanistic insight into how dual modulation of oestrogen feedback and aromatase inhibition can restore ovarian function more-effectively than monotherapy. The findings underscore the translational relevance of this combination approach as a potential strategy for clomiphene-resistant PCOS or patients with concurrent metabolic dysfunctions, thereby offering a foundation for future clinical trials and individualised therapeutic regimen.

## Funding

None

## Availability of data and materials

Data generated during and analysed during the course of this study are available from the corresponding author on request.

## Declaration of ethical approval

Ethical approval for the research was granted by the Ethical Committee of Faculty of Basic Clinical Sciences, LAUTECH.

## Competing interests

All authors of this paper declare that there is no conflict of interest related to the content of this manuscript.

## Notes

### Competing Interest Statement

The authors have declared no competing interest.

